# Kinetic investigation of calcium-induced Sorcin aggregation by stopped-flow light scattering

**DOI:** 10.1101/2025.03.03.641334

**Authors:** Qiushi Ye, Kathleen Joyce Carillo, Nicolas Delaeter, Lei Zhang, Jaekyun Jeon, Yanxin Liu

## Abstract

Sorcin, a penta-EF hand calcium-binding protein, is implicated in multidrug resistance (MDR) in various cancers and has roles in neurodegenerative diseases. It regulates cellular calcium homeostasis by interacting with calcium channels, pumps, and exchangers in a calcium-dependent manner. Calcium binding induces a conformational change in Sorcin, exposing hydrophobic surfaces that mediate protein interactions and calcium flux between the cytosol and endoplasmic reticulum (ER). These exposed surfaces can also drive Sorcin aggregation in the absence of binding partners. Here, we exploited calcium-induced conformational changes and aggregation of Sorcin as a model to study its calcium sensitivity and aggregation mechanisms. Stopped-flow light scattering revealed that Sorcin aggregation is reversible, cooperative, and primarily influenced by Sorcin concentration rather than physiological calcium levels. Our findings suggest that calcium sensitivity of Sorcin is finely tuned by its expression level, highlighting its role as an intracellular calcium sensor. This work establishes Sorcin as a model system for studying protein aggregation mechanisms with implications for MDR and neurodegenerative diseases.

## Introduction

Sorcin was first identified as a **<u>so</u>**luble **<u>r</u>**esistance-related **<u>c</u>**alcium-binding prote**<u>in</u>** in the K562 leukemia cell line^1^. Subsequently, it was recognized as an overexpressed protein in several human tumors exhibiting multidrug resistance (MDR), including gastric, breast, and ovarian cancers, as well as glioblastoma^2–9^. The Sorcin gene resides in the same chromosomal region and amplicon as MDR-related genes, such as ABC transporters MRD1 (ABCB1) and MRD3 (ABCB4), highlighting Sorcin’s role in chemotherapy resistance^10^. Furthermore, a mitochondrial isoform of Sorcin interacts with the mitochondrial chaperone Trap1^11^. These proteins are co-upregulated in colorectal carcinomas and jointly protect cells against apoptosis induced by chemotherapeutic agents^11^. Beyond its role in cancer, Sorcin is overexpressed in brain tissue from Alzheimer’s and Parkinson’s patients, where it may counteract the elevated cytosolic calcium (Ca^2+^) concentrations associated with neurodegeneration by facilitating Ca^2+^ uptake into the endoplasmic reticulum^12^. Due to its roles in both cancer and neurodegeneration, Sorcin has emerged as a promising drug target for therapeutic intervention^13,14^.

Beyond its involvement in human diseases, Sorcin plays a key role in regulating cellular Ca^2+^ homeostasis^15,16^. Upon Ca^2+^ binding, it undergoes a major conformational change that exposes hydrophobic surfaces, enhancing its interactions with other proteins^17–20^. In a Ca^2+^-dependent manner, Sorcin interacts with calcium channels, pumps and exchangers, including Ryanodine receptors (RyRs), Sarcoendoplasmic reticulum Ca^2+^ ATPase (SERCA), L-type voltage-dependent calcium channels, and Na^+^-Ca^2+^ exchangers (NCX)^21–25^. When cytosolic Ca^2+^ concentration is high, Sorcin inhibits Ca^2+^ release from the ER by interacting with RyRs while activating Ca^2+^ re-uptake through interactions with SERCA. This dual action promotes Ca^2+^ accumulation within the ER and reduces cytosolic Ca^2+^ levels, maintaining cellular calcium balance^26^.

Sorcin is a 21.6 kDa penta-EF hand calcium-binding protein and functions a homodimer^18^. Structurally, it consists of a glycine-rich N-terminal domain (NTD, residues 1-32) and a calcium-binding C-terminal domain (CBD, residues 33-198). X-ray crystallography and AlphaFold predictions show that the NTD is flexible in the absence of Ca^2+^ and may interact with the CBD upon Ca^2+^ binding^15,18^. Our recent structural characterizations of full-length Sorcin using NMR indicate that the NTD predominantly adopts an α-helical structure^27^. The CBD comprises five EF-hand motifs; while EF1-3 bind Ca^2+^ with varying affinities and contribute to Sorcin’s Ca^2+^ sensing, EF4 and EF5 are defective in Ca^2+^ binding, with EF5 playing a role in dimer formation. Upon Ca^2+^ binding, Sorcin undergoes a substantial conformational change, enabling it to bind and regulate target proteins in a calcium-dependent manner.

Here, we exploit Ca^2+^-induced Sorcin conformational change and resulting aggregation to investigate its Ca^2+^ sensitivity and aggregation mechanisms. Protein aggregation has predominantly been studied using fluorescence spectroscopy with aggregate-sensitive dyes, such as Thioflavin-T (ThT), or through turbidity assays that measure reduced transmitted light due to scattering. However, the limited temporal resolution of instruments like plate readers has made it challenging to capture rapid aggregation kinetics, particularly the early stages such as nucleation. The stopped-flow technique, widely used to monitor rapid chemical reactions, including protein folding on millisecond timescales, has significant potential for light scattering applications, though it had remained underexplored^28^. Recently, Jeon et al. demonstrated the use of stopped-flow light scattering to study the aggregation of amyloid-β (Aβ) peptides, providing quantitative analysis of Aβ oligomeric states over extended timescales, ranging from milliseconds to hours^29^.

Using stopped-flow light scattering, we observed that Sorcin aggregation is reversible and cooperative, with minimal sensitivity to physiological Ca^2+^ levels but high sensitivity to Sorcin concentration. Our results indicate that Sorcin’s Ca^2+^ sensitivity is finely tuned by its expression level, providing new insights into its role as an intracellular Ca^2+^ sensor and its contribution to MDR. These findings establish Sorcin as a robust model system for investigating protein aggregation mechanisms, with implications for both cancer resistance and neurodegenerative disease research.

## Methods

### Expression and purification of full-length human Sorcin

The DNA sequence encoding human Sorcin was codon-optimized for expression in *E. coli* and cloned into the pET151/D-TOPO vector, incorporating an N-terminal 6xHis tag followed by a TEV protease cleavage site. *E. coli* Rosetta (DE3) cells were transformed with this construct and grown in Lysogeny Broth (LB) with selective antibiotics. Cultures were incubated at 37°C until reaching an OD_600_ of 0.5–0.6, at which point they were cooled to 16°C and induced with 0.3 mM isopropyl-β-D-thiogalactoside (IPTG) for protein expression overnight. Cells were then harvested by centrifugation, resuspended in buffer, and lysed via sonication. The lysate was centrifuged to remove cellular debris, and the supernatant was purified using Ni^2+^-NTA affinity chromatography. The 6xHis tag was cleaved from Sorcin by TEV protease during overnight dialysis to remove imidazole. Further purification was achieved using anion-exchange chromatography (Mono Q column) and size-exclusion chromatography (Superdex 75 column). Protein purity was confirmed by SDS-PAGE, and concentration was determined by absorbance at 280 nm (ε = 25,900 M^−1^ cm^−1^).

### Turbidity assay

The turbidity assay was conducted at 25°C in a 384-well microplate using a SpectraMax iD5 plate reader to monitor protein aggregation via light scattering at 350 nm. The final reaction mixture consisted of 10 µM Sorcin, 100 µM CaCl_2_, 40 mM HEPES (pH 7.5, adjusted with KOH), 150 mM potassium chloride (KCl), and 1 mM β-mercaptoethanol (βME). To initiate the aggregation reaction, CaCl_2_ was added as the final component.

### Stopped-flow light scattering

Stopped-flow light scattering experiments were carried out on an Applied Photophysics SX20 spectrometer, with data collected using Applied Photophysics Pro-Data SX software. Solutions of Sorcin and CaCl_2_, prepared in matched buffer, were pre-loaded into two syringes. For each measurement, 60 µl from each syringe was pneumatically driven into the 120 µl optical chamber and rapidly mixed. A 565 nm LED served as the light source, and light scattering was detected at a 90° angle by a photomultiplier tube (PMT), with recorded voltage providing a measure of protein aggregation. Experiments were conducted at 22°C, with standard concentrations of 1 µM Sorcin and 150 µM CaCl_2_, unless otherwise specified in experiments. The buffer consisted of 40 mM HEPES (pH 7.5, adjusted with KOH), 150 mM KCl, and 1 mM βME, unless otherwise noted KCl titrations. Titration experiments were conducted for Sorcin (0.3 µM to 1.3 µM), Ca^2+^ (60 µM to 160 µM), Mg^2+^ (0 to 16 mM), and KCl (0 to 450 mM). All measurements were performed in triplicate.

### Kinetic analysis

The time series data obtained from stopped-flow light scattering were analyzed by fitting a double-exponential function to extract the kinetic parameters of Sorcin aggregation:

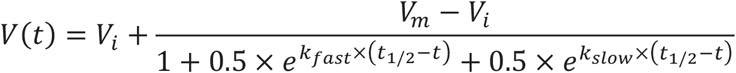

where V_i_ represents the initial background light scattering signal before aggregation begins, and V_max_ denotes the maximum light scattering signal, reflecting the final extent of aggregation. The parameter t_1/2_ is the aggregation half-time, at which the signal reaches the midpoint between V_i_ and V_m_. The rate constants k_fast_ and k_slow_ correspond to the fast and slow kinetic processes, respectively. The slope at t_1/2_ is defined as k_1/2_. The aggregation lag time (*τ*) was defined as the time at which V_i_ intersects with the tangent line, with slope k_1/2_, passing through the point (t_1/2_, (V_i_+V_m_)/2) on the kinetic curve. Mathematically, *τ* can be expressed as:

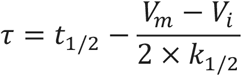

A double-exponential model was selected as it provided a significantly improved fit over a single-exponential model (Figure S1). For simplicity, equal weighting was applied to the two exponential terms, resulting in an excellent fit with minimal residuals.

The maximum aggregation rate (k_max_) and the time (η) at which k_max_ occurs were determined by calculating the first and second derivatives of the kinetic curves. Specifically, η represents the time at which the second derivative of the kinetic curve first reaches zero after the initial peak, marking the onset of the fastest aggregation rate. k_max_ is derived from the first derivative of the kinetic curve at time η. To minimize noise, a running average was applied to the kinetic data, with averaging intervals of 10 s for the aggregation data, 10 s for the first derivative, and 20 s for the second derivative. All derivative calculations were based on these smoothed data. Mean values and standard deviations for all kinetic parameters were obtained from three independent measurements.

## Results

### Ca^2+^-induced Sorcin aggregation is reversible

We confirmed that Ca^2+^ binding induces Sorcin aggregation, as previously reported^17^, by mixing Ca^2+^ with Sorcin in a test tube. Upon addition of Ca^2+^, the initially clear Sorcin solution visibly turned turbid, indicating aggregation (Figure 1B). When EGTA, a Ca^2+^ chelator, was subsequently added, the turbid Sorcin/Ca^2+^ mixture reverted to a clear solution, demonstrating that Sorcin aggregation is fully reversible upon Ca^2+^ removal through chelation by EGTA (Figure 1B). To monitor the aggregation process quantitatively, we utilized a turbidity assay, measuring light scattering at 350 nm. Aggregation kinetics were assessed using a plate reader (Figure 1C). While this approach captured the final equilibrium state, the initial aggregation phase occurred too rapidly to be resolved due to the plate reader’s limited time resolution (Figure 1C).

**Figure 1.**
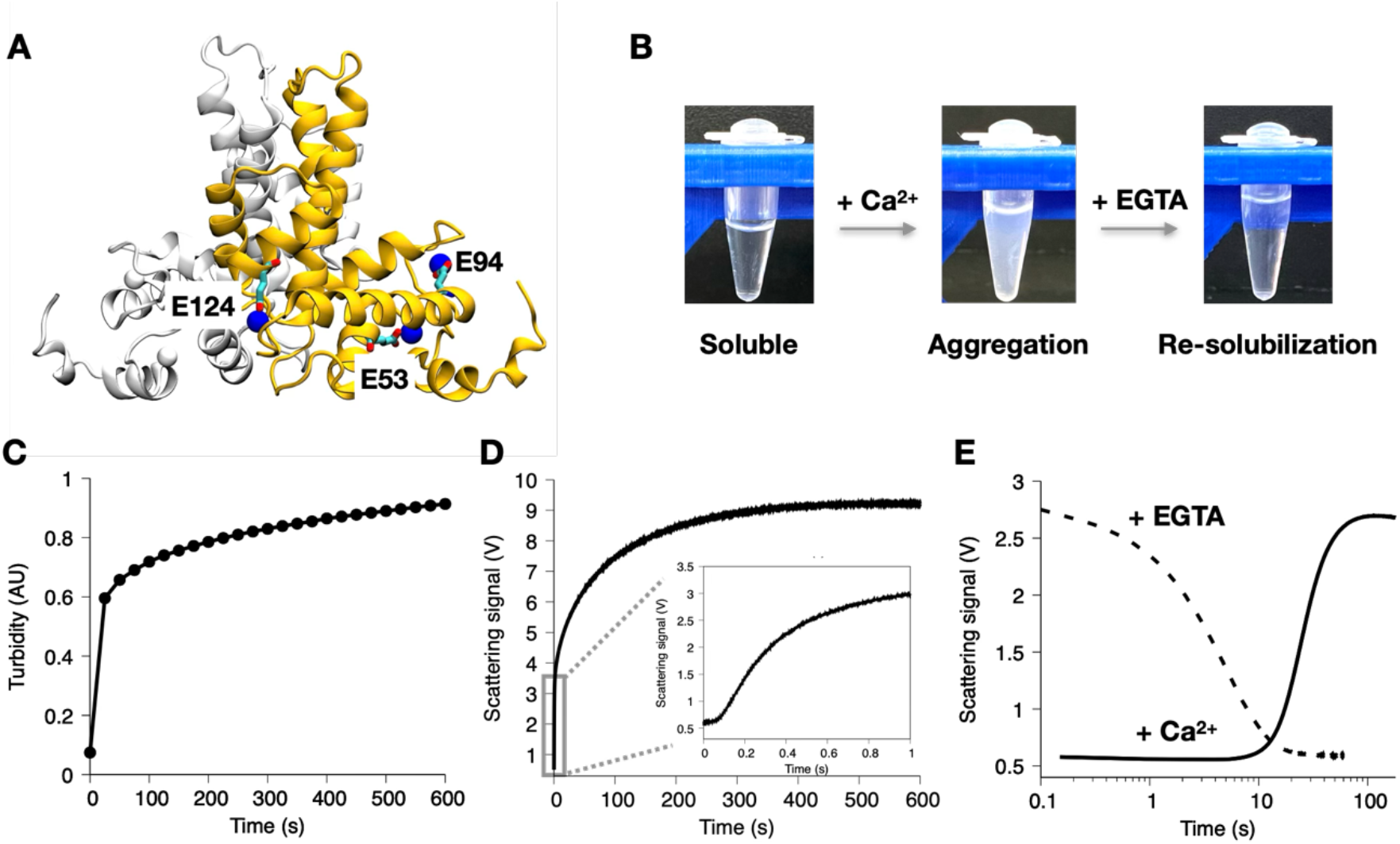
Calcium binding induces rapid and reversible Sorcin aggregation. (A) X-ray crystal structure of dimeric Sorcin in the Ca^2+^-bound state. The two protomers are shown in yellow and white, with key residues in the EF-hands labeled. Ca^2+^ ions are depicted in blue. (B) Ca^2+^ binding triggers Sorcin aggregation, which is reversed by the addition of EGTA. (C) Sorcin aggregation is monitored using a turbidity assay that measures light scattering at λ = 350 nm on a plate reader. (D) Stopped-flow light scattering experiments reveal fast initial aggregation kinetics with a ∼0.1 s lag phase. Light scattering signals were converted to voltage using a photomultiplier. The experiments were conducted by rapidly mixing 100 µM Ca^2+^ with 10 µM Sorcin, under the same conditions as the plate reader-based turbidity assay. (E) Reversal of Ca^2+^-induced Sorcin aggregation monitored by stopped flow light scattering. Aggregation of 1 µM Sorcin is induced by 150 µM Ca^2+^ with a 10 s lag phase (solid line) and reverses within 10 s upon adding 160 µM EGTA (dashed line).

To resolve these rapid kinetics, we employed stopped-flow light scattering, ideal for capturing reactions on the millisecond (ms) to second (s) timescale^29^. In our setup, a 565 nm LED illuminated the sample, and scattered light was detected by a photomultiplier tube (PMT) at a 90° angle. Aggregation was recorded as a change in voltage (V) detected by the PMT. Under the same conditions as the turbidity assay (10 µM Sorcin with 100 µM Ca^2+^), aggregation kinetics were recorded with a time resolution of 60 ms, as shown in Figure 1D. Consistent with turbidity assay results, aggregation reached a plateau within 10 minutes. Additionally, a lag phase of ∼0.1 seconds preceded the rapid growth phase, confirming that our stopped-flow instrumental setup is well-suited to study fast Sorcin aggregation kinetics.

With the stopped-flow light scattering implemented, we further investigated the re-solubilization rate of aggregated Sorcin upon EGTA addition. The highly sensitive PMT revealed that 1 µM Sorcin with 150 µM Ca^2+^ reached a plateau of 2.75 V with a 10-second lag (Figure 1E). Pre-incubated 1 µM Sorcin and 150 µM Ca^2+^ yielded the same signal of 2.75 V, but the addition of 160 µM EGTA rapidly dissolved the aggregates within 10 seconds (Figure 1E), confirming the reversibility of Ca^2+^-induced Sorcin aggregation and showing that re-solubilization occurs as quickly as aggregation.

### Effect of protein concentration on sorcin aggregation

Using stopped-flow light scattering, we observed that Sorcin aggregated at a concentration of 1 µM (Figure 1E), ten times lower than in our plate reader-based turbidity assays (Figure 1C). However, to induce aggregation at this low Sorcin concentration, a higher Ca^2+^ concentration (150 µM) was needed, resulting in a lag time approximately 100-fold longer than with 10 µM Sorcin. These results suggest that protein concentration influence Sorcin aggregation kinetics, a dependence commonly observed in protein aggregation.

To further explore protein concentration effects, we varied Sorcin concentrations from 0.3 µM to 1.3 µM in stopped-flow light scattering with a fixed 150 µM Ca^2+^ concentration. Representative aggregation traces are shown in Figure 2A. The first and second derivatives of the aggregation kinetics curves (Figures 2B and 2C) reveal the maximum aggregation rate (k_max_) and the time (η) at which it occurs, respectively. As expected, higher Sorcin concentrations led to increased aggregation, indicated by higher final scattering plateaus, and accelerated aggregation (Figure 2A). Fitting the data with a double-exponential function (see Methods), we found that aggregate levels, quantified by the maximum scattering signal (V_m_), increased linearly with Sorcin concentration, with a slope of 2.82 ± 0.08 V/µM (Figure 2D).

**Figure 2.**
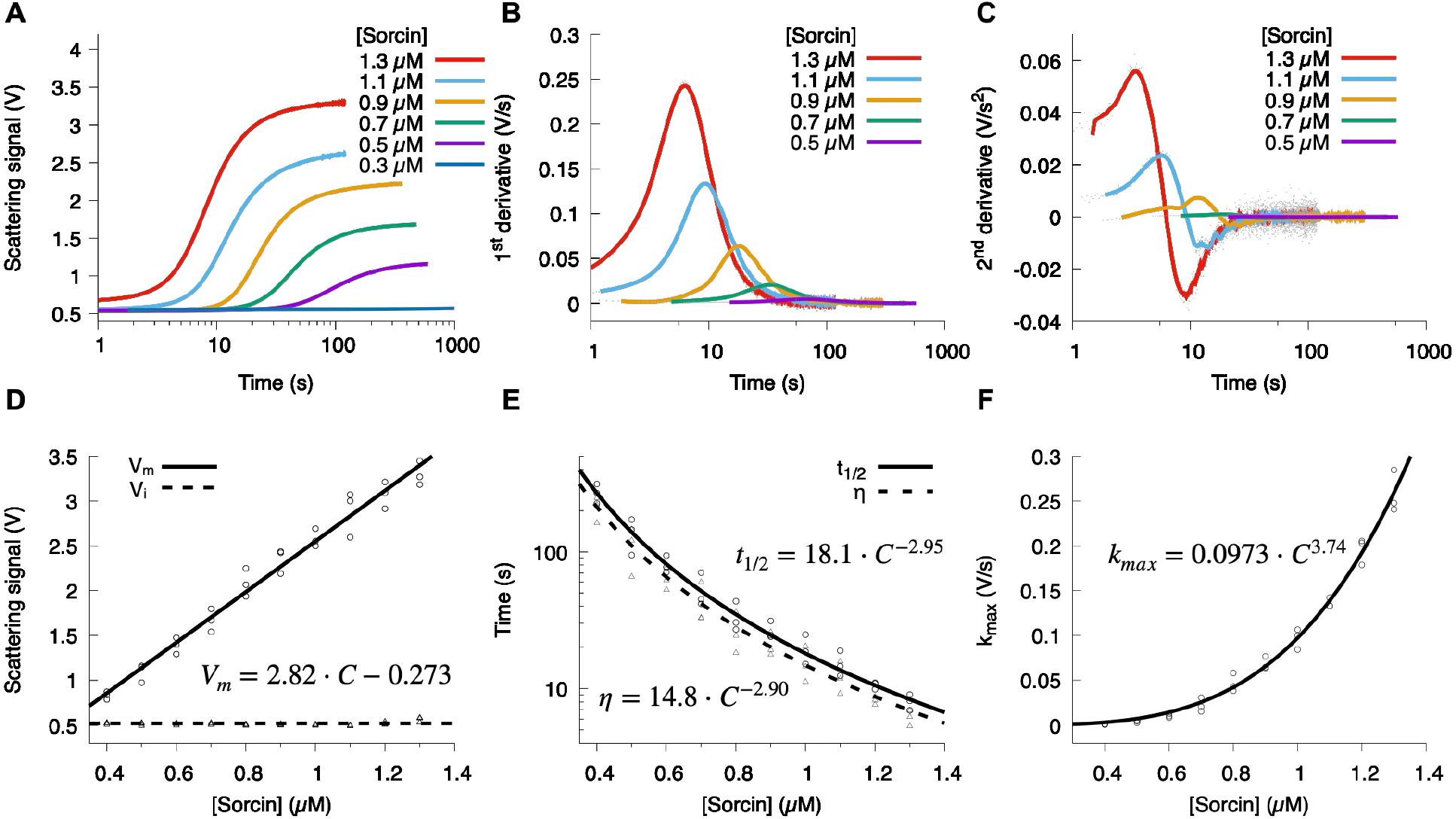
Sorcin aggregation kinetics depend on protein concentration. (A) Representative aggregation kinetics curves at varying Sorcin concentrations. (B) The first derivative of these curves is presented. (C) The second derivative, with raw data shown as gray dots and a running average (solid line) to reduce noise. (D) The signal plateau (V_m_), representing the maximum extent of aggregation, scales linearly with protein concentration, while initial signal V_i_ remains constant. Extracted V_m_ and V_i_ values are shown as circles and triangles, respectively, with linear and horizontal fits depicted as solid and dashed lines. (E) The aggregation half-time (t_1/2_) and the time at which the aggregation reaches its maximum rate (η) both follow a power law with a scaling exponent of ∼3. t_1/2_ and η values from individual experiments are shown as circles and triangles, and power-law fits as solid and dashed lines, respectively. (F) The maximum aggregation growth rates (k_max_) depend on Sorcin concentration and were fitted to a power-law model (solid line).

The double-exponential fit also yielded other kinetic parameters, including the aggregation half-time (t_1/2_) and two rate constants. As Sorcin concentration increased, t_1/2_ shortened according to a power law, *t*_1/2_= 18.1 ± 2.49 × *C*^(−2.95±0.17)^, indicating a complex aggregation mechanism potentially involving secondary nucleation or growth (Figure 2E). Both k_fast_ and k_slow_ also increased with Sorcin concentration (Figure S2).

We calculated the maximum aggregation rate (k_max_) and the time (η) at which it occurs by taking derivatives of the kinetics trace (Figures 2B and 2C). The dependence of η on Sorcin concentration followed a similar power law η = 14.9 ± 2.59 × *C*^(−2.90±0.21)^, aligning with the t_1/2_ exponent, validating our double-exponential fit approach (Figure 2E). The observed η was consistently shorter than t_1/2_ at each protein concentration, suggesting a fast initial process followed by slower growth, reinforcing the need for a double-exponential fit (Figure 2E). k_max_ also followed a power law *k*_*max*_ = 0.0973 ± 0.002 × *C*^(3.74±0.08)^ as protein concentration increased (Figure 2F), and higher Sorcin concentrations decrease lag time *τ* (Figure S2).

### Effect of Ca^2+^ concentration on sorcin aggregation

Given that Sorcin functions as a Ca^2+^ sensor, we expected that aggregation would depend on Ca^2+^ concentration. Indeed, higher Ca^2+^ concentrations led to more extensive and faster aggregation as shown in Figure 3A. Unlike the linear dependence of V_m_ on Sorcin concentration, the V_m_ response to Ca^2+^ concentration was sigmoidal, best fit by a Hill equation, 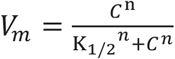 as shown in Figure 3B. A Hill coefficient of n=9.15 ± 0.89 indicates high cooperativity in Ca^2+^-induced aggregation, with half of Sorcin aggregating at K_1/2_=93.7 ± 1.10 µM Ca^2+^. Higher Ca^2+^ concentrations also shortened all aggregation timescales, including t_1/2_, η, and τ, and accelerated all kinetic rates (Figure S3).

**Figure 3.**
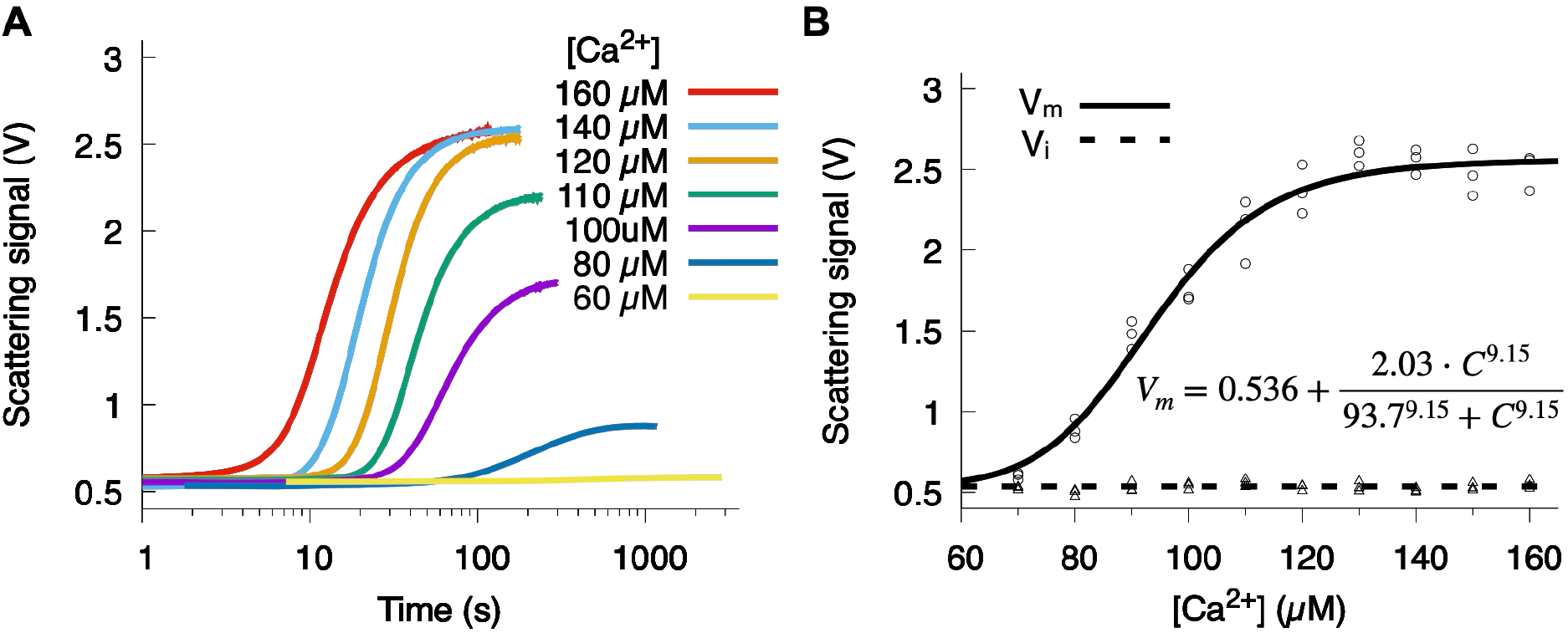
Ca^2+^ concentration dependence of the sorcin aggregation kinetics. (A) Representative aggregation kinetics curves at varying Ca^2+^ concentrations. (B) The dependence of V_max_ on Ca^2+^ concentration follows a sigmoidal response. V_max_ and V_min_ values from individual experiments are shown as circles and triangles, respectively. The relationship between V_max_ and Ca^2+^ concentration is fitted with a Hill equation (solid line), while V_min_, which remains constant across different Ca^2+^ concentrations, is fitted with a horizontal line (dashed line).

### Mg^2+^ inhibition of Ca^2+^-induced Sorcin aggregation

Sorcin can also bind Mg^2+^, which is present at higher cellular concentrations than Ca^2+^ in resting states. We examined how physiological Mg^2+^ levels affect Ca^2+^-induced Sorcin aggregation by repeating stopped-flow light scattering experiments (1 µM Sorcin with 150 µM Ca^2+^) in the presence of Mg^2+^. At < 2 mM Mg^2+^, there was no effect on aggregation, but concentrations >4 mM significantly inhibited Ca^2+^-induced aggregation (Figures 4A-4C). The aggregation rates were shown in Figure S4. These findings suggest that Mg^2+^ competes with Ca^2+^ for binding to Sorcin’s EF hands, modulating Sorcin sensitivity to Ca^2+^ in a manner dependent on Mg^2+^ concentration, which varies across cell types and states.

**Figure 4.**
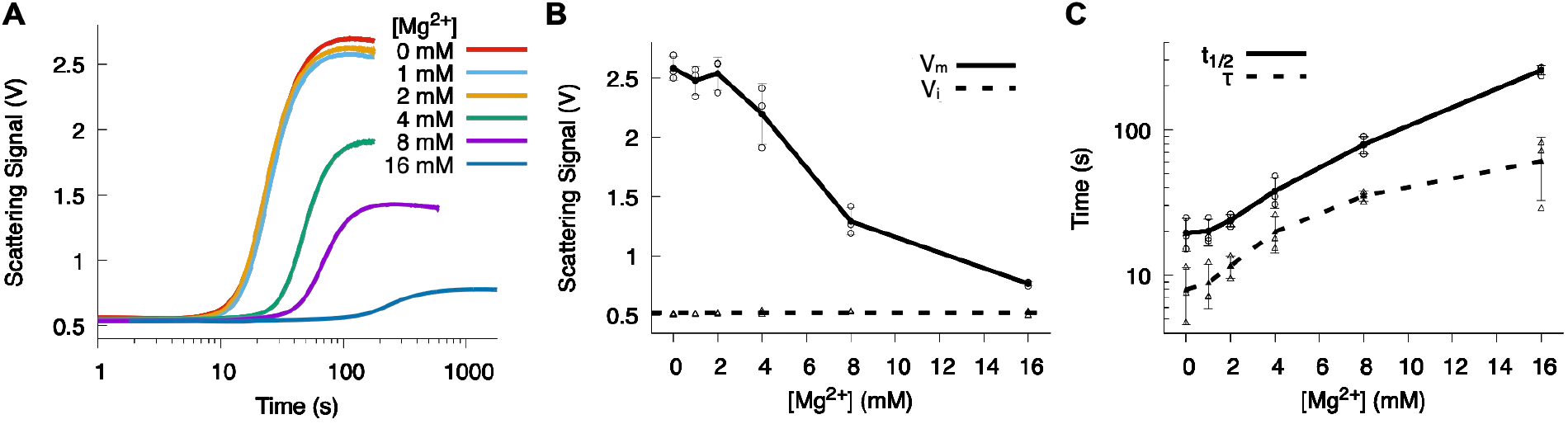
Effect of Mg^2+^ on Sorcin aggregation. (A) Representative aggregation kinetics curves at varying Mg^2+^concentrations. (B) The dependence of V_max_ on Mg^2+^ concentration, with the average V_max_ from three independent experiments at each Mg^2+^ concentration connected by a solid line. (C) The dependence of the aggregation half-time (t_1/2_) and lag time (τ) on Mg^2+^concentration.

### Effect of salt concentration on Sorcin aggregation

Salt concentration can impact protein aggregation by screening charges and weakening protein-protein interactions. To test this effect, we varied KCl concentration from 2 mM to 430 mM. High salt levels inhibited aggregation, as reflected by lower V_m_ (Figure 5A and 5B), increased t_1/2_ (Figure 5C), and slower rates (Figure S4). Surprisingly, very low salt concentrations (<30 mM) also inhibited aggregation, potentially by stabilizing intramolecular interactions in the Ca^2+^-free state of Sorcin. This stabilization could increase the energy barrier for transitioning to the Ca^2+^-bound, aggregation-prone state. Further studies are needed to elucidate the molecular mechanisms behind this unusual salt dependence.

**Figure 5.**
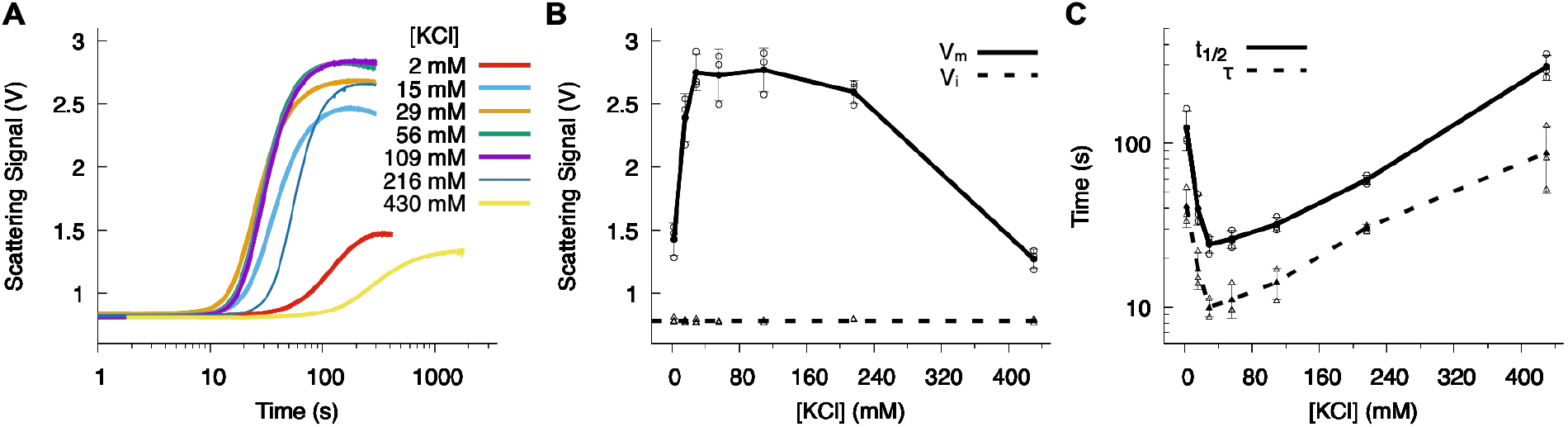
Effect of salt concentrations on Sorcin aggregation. (A) Representative aggregation kinetics curves at varying KCl concentrations. (B) The dependence of V_max_ on KCl concentration, with the average V_max_ from three independent experiments at each KCl concentration connected by a solid line. (C) The dependence of the aggregation half-time (t_1/2_) and lag time (τ) on KCl concentration.

### Effect of EF-hand mutations on Sorcin aggregation

Sorcin is a penta-EF-hand protein, but only the first three EF hands bind Ca^2+^ due to sequence mutations in EF4 and EF5^30^. We tested point mutations in EF1-3 (E53Q, E94A, and E124A) to assess their impact on Ca^2+^-induced aggregation. Each mutation was sufficient to abolish aggregation at basal conditions (1 µM Sorcin and 150 µM Ca^2+^) (Figure S5), confirming their effectiveness in disrupting Ca^2+^ binding.

Increasing protein or Ca^2+^ concentrations partially restored aggregation for specific mutants as shown in Figure 6. At 1.3 µM Sorcin and 150 µM Ca^2+^, wild-type Sorcin aggregated rapidly (V_m_ = 3.3 V, t_1/2_= 8 s), while none of the mutants aggregated. At 1 mM Ca^2+^, the EF2 mutant (E94A) reached a V_m_ similar to wild-type with a slower aggregation kinetics (t_1/2_= 14 s), and EF1 mutant (E53Q) aggregated less and even slower (V_m_ = 0.7 V, t_1/2_ = 264 s). EF3 mutant (E124A) required 2 µM protein and 2 mM Ca^2+^ to show mild aggregation (V_m_ = 0.6 V, t_1/2_ = 204 s). These findings suggest cooperative interactions among EF hands, with each EF hand contributing differentially to Ca^2+^-sensitivity and aggregation.

**Figure 6.**
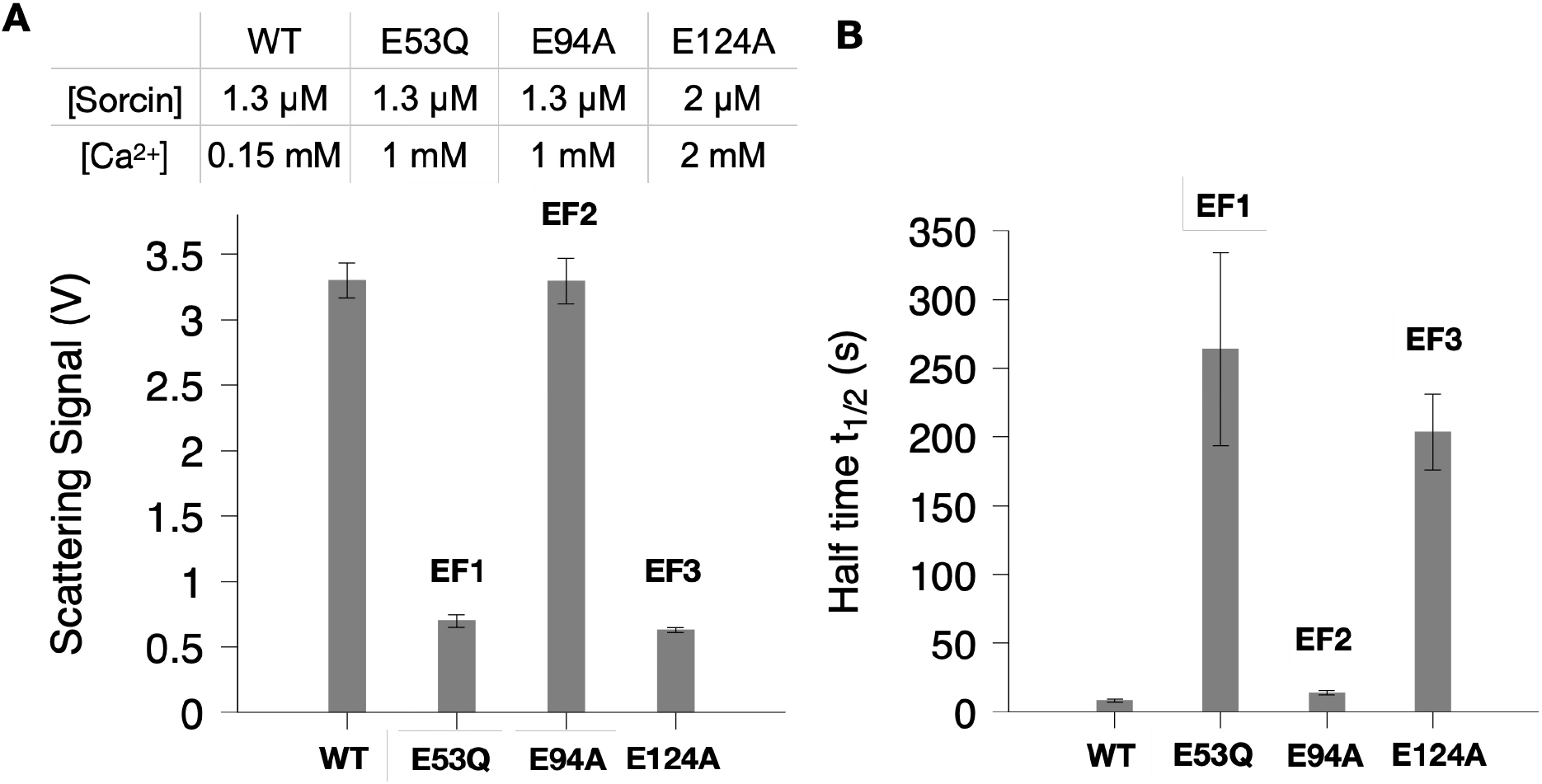
Effect of mutations in three EF hands on Sorcin aggregation. (A) Effect of the mutations on V_max_. (B) Effect of the mutations on the aggregation half-time (t_1/2_). Note that all aggregation assays for the mutations were performed at either higher Sorcin or Ca^2+^ concentrations compared to the wild-type experiments, as indicated in the table. No aggregation was observed for the mutations under the conditions used for the wild-type aggregation experiments (see Figure S5).

## Discussions

A major function of Sorcin is to regulate intracellular Ca^2+^ levels by interacting with Ca^2+^ channels and pumps on the endoplasmic reticulum (ER) membrane^21–25^. This regulation is initiated by Sorcin’s ability to sense changes in Ca^2+^ concentration: when Ca^2+^ levels rise, Sorcin binds Ca^2+^ via its EF-hand motifs, triggering a conformational change that exposes hydrophobic residues on its surface^15^. This exposed hydrophobic region mediates Sorcin’s interactions with Ca^2+^ channels and pumps, ultimately suppressing Ca^2+^ release from channels and activating pumps to sequester excess cytosolic Ca^2+^ back into the ER^31^. In the absence of binding partners, Ca^2+^-bound Sorcin self-associates through these hydrophobic surfaces, leading to protein aggregation in vitro^18^. Here, we implemented stopped-flow light scattering to capture the rapid kinetics of Ca^2+^-induced Sorcin aggregation.

A unique feature distinguishing Sorcin from other aggregation-prone proteins like Aβ peptides and α-synuclein is its reversible aggregation^17,32,33^. Ca^2+^-induced Sorcin aggregation was fully reversible upon Ca^2+^ removal with EGTA, as demonstrated in Figure 1. Remarkably, the disaggregation rate was as fast as the aggregation process, suggesting that Sorcin aggregates form via interactions between well-folded proteins rather than by unfolded or misfolded proteins. This characteristic contrasts with most well-studied aggregation systems, where aggregation is often linked to misfolding or alternative structural states. Sorcin thus provides a unique system for studying reversible aggregation kinetics and thermodynamics, offering insight relevant to understanding aggregation in pathological contexts.

As a cellular Ca^2+^ sensor, a key question about Sorcin is its sensitivity to Ca^2+^ concentrations. Although EF-hands 1 and 2 bind Ca^2+^ with μM affinity, no Sorcin aggregation was observed at Ca^2+^ concentrations in the μM range. This suggests that either Sorcin conformational change is not triggered by Ca^2+^ binding to EF-hands 1 and 2, or that Sorcin responds to μM Ca^2+^ concentrations through a mechanism other than hydrophobic surface exposure. Aggregation was only observed at Ca^2+^ concentrations above 60 μM, with half-maximal aggregation at 93 μM, suggesting that EF-hand 3 may have a Ca^2+^ affinity in the tens of μM range (Figure 3). The result renders Sorcin as a much less sensitive Ca^2+^ than well-studied Ca^2+^ binding protein Calmodulin which has sub-micromolar and micromolar Ca^2+^ affinity^34^. Moreover, the presence of mM levels of Mg^2+^, typical in cellular environments, may increase Ca^2+^ binding affinity to the hundreds of μM range (Figure 4A-4C). Given that resting cytosolic Ca^2+^ levels are generally around 100 nM, Sorcin may be primarily responsive in specific cellular contexts, such as in certain cell types or developmental stages where Ca^2+^ concentrations temporarily reach elevated levels (e.g., in excitable cells such as cardiomyocytes during stimulation).

While Sorcin aggregation shows limited sensitivity to Ca^2+^, it is highly responsive to Sorcin protein concentration. Increasing Sorcin levels from 0.4 μM to 1.3 μM decreased the aggregation half-time by an order of magnitude, from ∼300 s to ∼10 s (Figure 2). This high sensitivity to Sorcin concentration may allow cells to fine-tune Ca^2+^ responses by modulating Sorcin expression. Additionally, this responsiveness appears to be optimized at physiological salt concentrations (∼100 mM). Deviations from this salt concentration significantly slowed aggregation, suggesting that Ca^2+^, sensitivity of Sorcin may also be influenced by ionic strength (Figure 5A-5C).

Beyond elucidating Sorcin’s role in Ca^2+^ sensing and regulation, our study highlights Sorcin as a model system for understanding protein aggregation mechanisms. Unlike classical aggregation systems such as amyloid-β, α-synuclein, tau, Huntington protein with polyglutamine extensions, and prion protein—where aggregation studies typically require extended incubation under shaking or elevated temperatures—Sorcin aggregates rapidly and reversibly under easily controlled conditions. These features make Sorcin an ideal system for studying aggregation kinetics. Furthermore, Sorcin’s aggregation is highly tunable by protein, Ca^2+^, Mg^2+^, and ionic strength. The complex and cooperative nature of Sorcin aggregation revealed by stopped-flow light scattering provides a framework for investigating the fundamentals of protein aggregation, a process linked to various neurodegenerative diseases. Recent findings of Sorcin as an early biomarker in neurodegenerative conditions underscore the need for further mechanistic insights into its aggregation behavior^12^.

## Conclusion

We implemented stopped-flow light scattering to investigate Sorcin’s rapid calcium-induced aggregation kinetics. Our results reveal that Sorcin aggregation is reversible, cooperative, and primarily driven by protein concentration rather than physiological calcium levels. These findings highlight the critical role of Sorcin expression in modulating its calcium sensitivity and aggregation behavior. This study advances our understanding of Sorcin’s function in calcium homeostasis and establishes it as a valuable model system for exploring protein aggregation mechanisms, with potential relevance to MDR and neurodegenerative diseases.

## Notes

The authors declare no competing financial interests.

## Acknowledgements

The authors thank the supports from all lab members in the Liu Lab at the University of Maryland College Park (UMCP). The project is supported by start-up funding from the University of Maryland College Park and the Institute for Bioscience and Biotechnology Research Seed Grant funded by the University of Maryland Strategic Partnership: MPowering the State.

