## Supporting Information for "Kinetic investigation of calcium-induced Sorcin aggregation by stopped-flow light scattering"

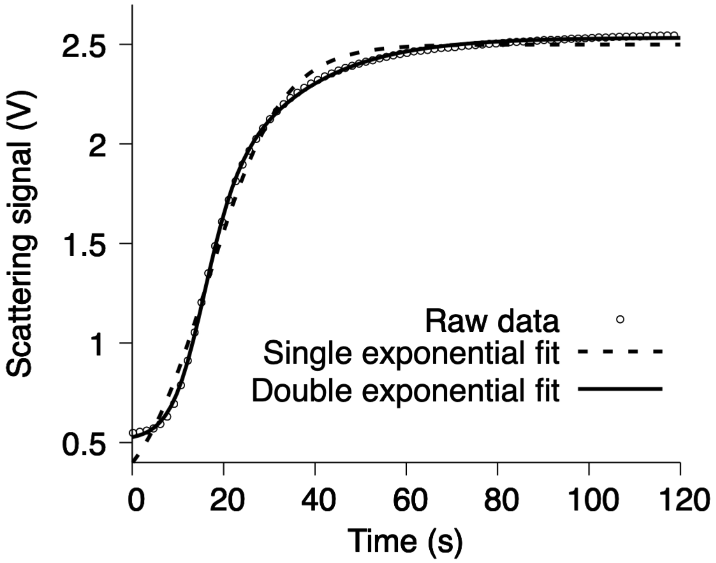


**Figure S1. Comparison between single and double exponential fits to Sorcin aggregation kinetics.** The light scattering kinetics trace (circles) was obtained by rapidly mixing 1 μM Sorcin with 150 μM $Ca^{2+}$ using stopped flow. The dashed line represents the single-exponential fit, yielding a root mean square deviation (RMSD) of 0.047 between the raw data and the fit. The solid line represents the double-exponential fit, yielding a significantly lower RMSD of 0.009, indicating a better fit to the data.

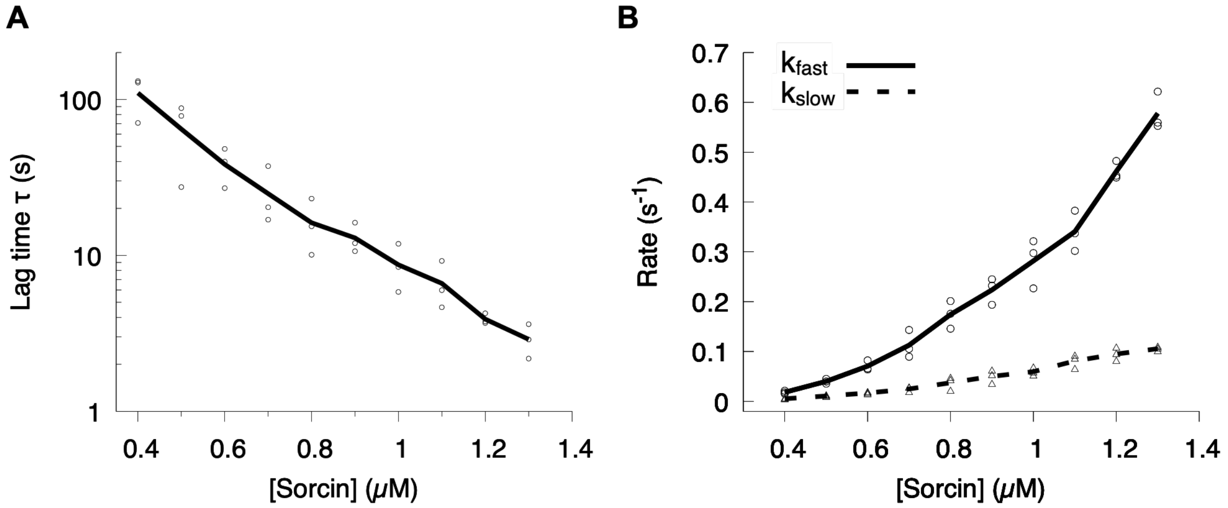


**Figure S2. Dependence of the aggregation lag time τ and the aggregation growth rates ($k_{fast}$ and $_{kslow}$) on protein concentration.** (A) The τ values calculated from three independent experiments at each Sorcin concentration are shown as circles, and the average τ at each concentration is connected by a solid line. (B) The aggregation growth rates ($k_{fast}$, and $_{kslow}$) depend on Sorcin concentration.

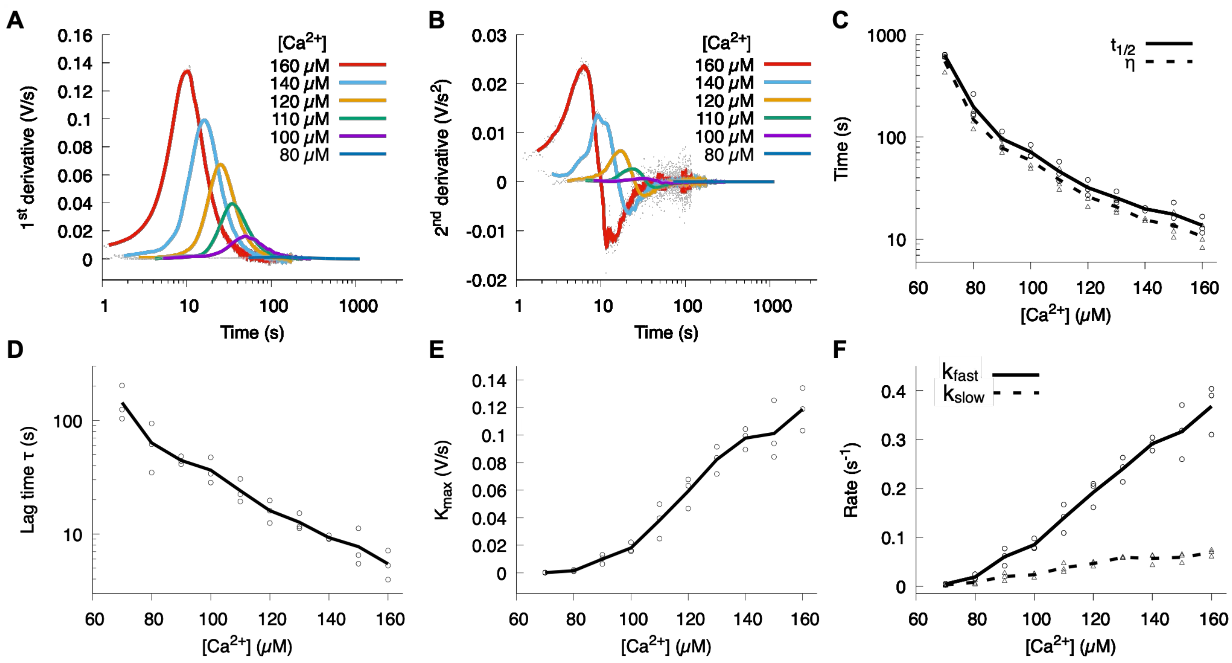


**Figure S3. Sorcin aggregation kinetics depend on $Ca^{2+}$ concentration.** (A) First derivative of representative aggregation kinetics curves at varying $Ca^{2+}$ concentrations. (B) Second derivative of these curves, with raw data shown as gray dots and a running average (solid line) applied to reduce noise. (C) Both the aggregation half-time ($t_{1/2}$) and the time at which aggregation reaches its maximum rate (η) depend on $Ca^{2+}$ concentration. Values extracted from individual experiments are shown as circles ($t_{1/2}$) and triangles (η) with averages connected by solid and dash lines, respectively. (D) The lag time (τ) depends on $Ca^{2+}$ concentration, and the value represents by circle with average connected by solid line. (E) The maximum aggregation growth rate ($k_{max}$) depends on $Ca^{2+}$ concentration. (F) The aggregation growth rates ($k_{fast}$ and $k_{slow}$) depend on $Ca^{2+}$ concentration.

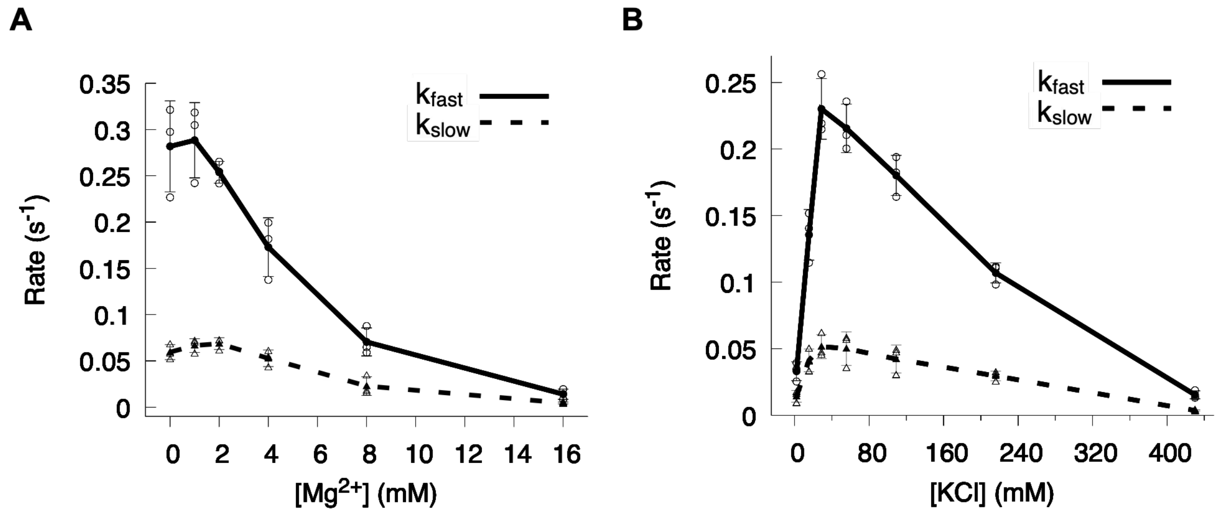


**Figure S4. Effect of $Mg^{2+}$ and salt concentrations on the rate of Sorcin aggregation.** (A) The Sorcin aggregation growth rates ($k_{fast}$ and $k_{slow}$) decrease as $Mg^{2+}$ concentration increases. Values from individual experiments are shown as circles ($k_{fast}$) and triangles ($k_{slow}$), with averages connected by lines. (B) Dependence of Sorcin aggregation growth rates ($k_{fast}$ and $k_{slow}$) on KCl concentration. Values from individual experiments are shown as circles ($k_{fast}$) and triangles ($k_{slow}$), with averages connected by lines.

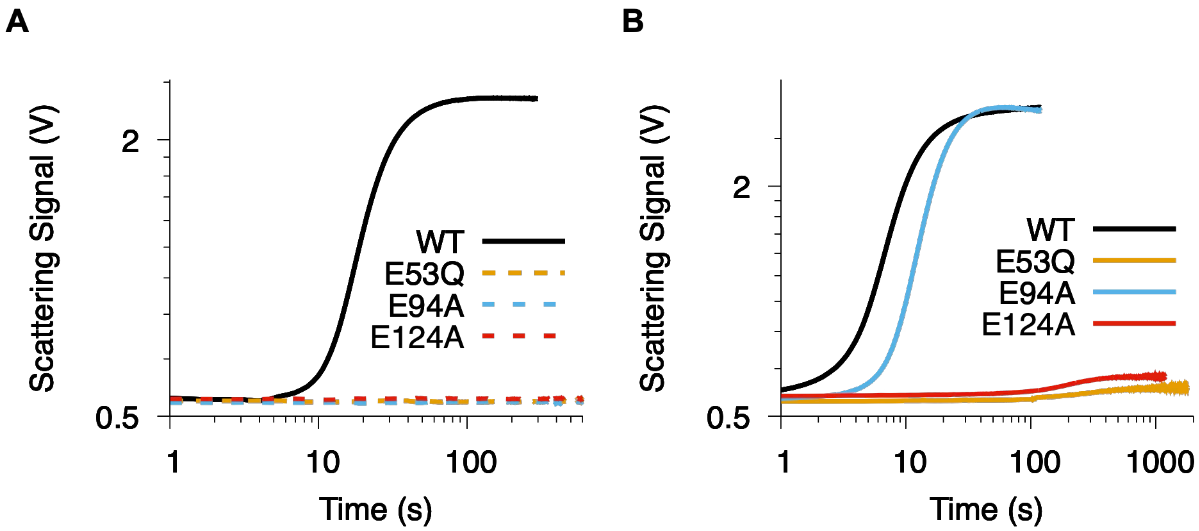


**Figure 5. Effect of mutations in three EF hands on Sorcin aggregation.** (A) Representative Sorcin aggregation kinetics curves for 1 µM Sorcin (wildtype or EF hand mutants) mixed with 150 µM $Ca^{2+}$. No aggregation was observed for the EF hand mutants under these conditions. (B) Representative Sorcin aggregation kinetics curves at increased Sorcin and $Ca^{2+}$ concentrations. The specific conditions are as follows: wildtype Sorcin (1.3 µM Sorcin, 150 µM $Ca^{2+}$), E53Q and E94A mutants (1.3 µM Sorcin, 1 mM $Ca^{2+}$) and E124A mutant (2 µM Sorcin, 2 mM $Ca^{2+}$).
